# Neurophysiological Effects of Sudarshan Kriya Revealed by Subject-Independent EEG Machine Learning

**DOI:** 10.64898/2026.03.09.710276

**Authors:** Abhishek Krishna Pillai, Oleksandr Bezrukavyi, Shaija Nalinakshan

**Author notes:** Corresponding Author: Abhishek Krishna Pillai Department of Data Analytics, Dublin Business School, Dublin, Ireland.

## Abstract

Sudarshan Kriya Yoga (SKY) is a structured breathing-based meditation associated with improvements in stress regulation and mental health, yet its acute neurophysiological effects remain incompletely characterized using multivariate, subject-independent approaches. In this study, we investigated short-term pre–post neural modulation induced by a single long SKY session using electroencephalography (EEG) and traditional machine learning. EEG signals were pre-processed, and multiple feature representations were extracted, including raw EEG voltage statistics (mean, standard deviation), Short-Time Fourier Transform (STFT)–based spectral features, Discrete Wavelet Transform (DWT)–based multi-resolution features, and coherence-based functional connectivity measures. Classification was performed under a leakage-free Leave-One-Subject-Out Cross-Validation (LOSO-CV) framework to assess subject-level generalizability.

Across feature types and classifiers, the intervention group consistently showed higher accuracy and more stable performance than the control group, indicating systematic SKY-related neural changes. STFT-based spectral features achieved the highest peak performance, with a Multilayer Perceptron reaching ∼89% accuracy, followed by DWT and coherence-based features, which also exhibited strong discriminative capacity. Raw EEG voltage features yielded lower performance, suggesting limited specificity. Control group classification remained close to the 0.50 baseline expected under random classification across all feature types, supporting the specificity of intervention effects.

These results demonstrate that acute SKY practice induces measurable neural modulation most robustly captured in frequency-domain and network-level EEG representations. The combination of subject-independent validation and comparative feature analysis provides a rigorous, data-driven framework for characterizing meditation-related brain dynamics and advancing the objective quantification of breathing-based interventions.

## 1. Introduction

Mental and neurological disorders constitute a growing global public health challenge in the modern era, despite unprecedented technological advancement, economic globalisation, and societal connectivity. Psychological distress—encompassing chronic stress, anxiety, and depressive disorders—has become increasingly prevalent worldwide, contributing substantially to disability, reduced quality of life, and economic burden. Epidemiological evidence indicates that common mental disorders, particularly depression and anxiety, continue to rise globally and represent a major challenge for health systems and societies at large [1].

At the core of this escalating burden lies chronic, unmanaged stress. Stress, classically defined as the body’s non-specific response to any demand placed upon it, is an adaptive mechanism essential for survival and performance under acute conditions [2]. However, when stress exposure is prolonged, sustained activation of physiological stress-response systems becomes maladaptive, leading to allostatic overload. Persistent psychosocial stressors—such as occupational demands, financial insecurity, social isolation, and rapid societal change—can maintain a state of chronic hyper-arousal, progressively depleting adaptive capacity and increasing susceptibility to both mental and physical illness [3,4].

The neurobiological foundation of the stress response is primarily governed by the hypothalamic–pituitary–adrenal (HPA) axis. In response to perceived threats, the hypothalamus releases corticotropin-releasing hormone, initiating a hormonal cascade that culminates in the secretion of cortisol from the adrenal glands. While acute cortisol release supports energy mobilisation and cognitive focus, prolonged hypercortisolemia leads to allostatic overload, exerting deleterious effects across multiple physiological systems [3]. Chronic stress exposure has been associated with structural and functional alterations in the central nervous system, including hippocampal volume reduction, amygdalar hyperactivity, and impaired prefrontal cortical connectivity—neural changes strongly implicated in the pathophysiology of depressive and anxiety disorders [4].

Chronic stress also disrupts autonomic nervous system regulation, leading to sustained sympathetic dominance and reduced parasympathetic activity. This imbalance contributes to both psychological distress and adverse physical health outcomes, including cardiovascular and metabolic dysfunction [5]. Consistent with this, heart rate variability—a non-invasive index of autonomic function—is reliably reduced in individuals with chronic stress, anxiety, and depression, reflecting impaired parasympathetic regulation and reduced physiological flexibility [6].

Despite advances in pharmacological and psychotherapeutic treatments, important limitations persist, including adverse effects, limited accessibility and affordability, challenges with long-term adherence, and insufficient impact on underlying physiological stress dysregulation. These challenges have spurred growing interest in accessible and scalable non-pharmacological interventions that target both psychological symptoms and their physiological mechanisms. Mind–body practices rooted in yoga and meditation have gained attention for their capacity to modulate stress responses through controlled breathing and attentional regulation, influencing autonomic balance, HPA axis activity, and central nervous system function, thereby enhancing emotional regulation and stress resilience [7–10].

Within this context, Sudarshan Kriya Yoga (SKY) has emerged as a structured, breathing-based meditation practice with demonstrated benefits across a range of psychological and physiological domains. Clinical and experimental studies have reported reductions in stress, anxiety, depressive symptoms, and burnout following SKY practice, alongside improvements in autonomic and neurophysiological markers [11–14]. However, despite growing evidence of its therapeutic potential, objective and methodologically rigorous characterization of SKY-induced neural changes remains limited.

SKY practice comprises different session formats, including a short kriya protocol of approximately 30 minutes and a long kriya protocol of approximately 45 minutes. While several prior EEG studies have examined the effects of sustained SKY practice over weeks or months, the immediate neurophysiological effects of a single session of long SKY have not been systematically examined. Moreover, existing studies have largely relied on conventional univariate statistical analyses, limiting assessment of subject-level generalizability.

In addition, although a small number of studies have applied machine learning techniques to EEG data in the context of SKY, key methodological components—such as rigorous preprocessing pipelines and subject-independent evaluation using Leave-One-Subject-Out Cross-Validation (LOSO-CV)—have not been systematically examined.

The present study addresses these gaps by evaluating whether traditional machine learning models can reliably distinguish pre- and post-intervention EEG activity following a single session of long Sudarshan Kriya, using time-domain, time–frequency, and connectivity-based features within a leakage-free LOSO-CV framework.

### Research Question

Can traditional machine learning models reliably distinguish pre- and post-intervention EEG activity associated with SKY using time-domain, time–frequency, and connectivity-based features?

### Hypothesis

Post-intervention EEG recordings will exhibit distinct neural patterns compared to pre-intervention recordings, enabling reliable classification using traditional machine learning models.

## 2. Methods

### Study Design and Participants

This study employed a within-subject experimental design to investigate neural changes associated with SKY. All EEG data used in the present analysis were obtained from a previously published study [15], and no new data were collected. EEG recordings were obtained from an intervention group consisting of experienced SKY practitioners (N = 40) and an active control group (N = 10). The control group participated in a guided music-based relaxation session, selected to provide a comparative intervention involving auditory stimulation and relaxation without structured breathing or meditative techniques.

For participants in the intervention group, EEG data were recorded under two conditions—pre-SKY and post-SKY—allowing each participant to serve as their own control. This design ensured equal representation of pre- and post-intervention samples at the subject level and minimized inter-individual variability. As each participant contributed paired recordings across conditions, no class imbalance was present.

Participants in the control group underwent EEG recording during a comparable pre- and post-session interval surrounding the music-based relaxation condition. Control group data were analyzed independently and were not paired with the SKY intervention data. The inclusion of a relaxation-based active control was intended to mitigate expectancy effects and control for non-specific influences such as attentional engagement and general relaxation.

All participant recruitment, experimental procedures, and ethical approvals were conducted as part of the original study [15]. The present work involved only secondary analysis of anonymized EEG data.

Demographic characteristics of the participants were reported in the original study [15]. Briefly, the intervention group consisted of 40 SKY practitioners (25 males, 15 females; mean age = 24.9 ± 5.5 years; meditation experience = 4.3 ± 3.9 years), while the control group included 10 participants (4 males, 6 females; mean age = 32.4 ± 11.7 years). Three additional intervention participants were excluded in the original study due to poor signal quality.

### EEG Data Acquisition and Preprocessing

Continuous EEG signals were recorded using a 32-channel system positioned according to the international 10–20 electrode placement standard. Acquisition parameters (sampling rate, referencing scheme, recording environment, and session timing) followed the procedures described in the original study [15]. Data were acquired during resting-state conditions immediately before and after the SKY intervention for the intervention group, and under equivalent resting conditions for the control group [15].

EEG preprocessing was performed to enhance signal quality and reduce contamination from non-neural sources. The preprocessing pipeline included:

1. **Bandpass filtering** to retain physiologically relevant EEG frequencies (e.g., 0.5–50 Hz).
2. **Notch filtering** to suppress power-line interference.
3. **Segmentation** of the pre-processed EEG signals into fixed-length epochs for feature extraction.

To ensure consistency across conditions and participants, equal-duration EEG segments were retained for both pre- and post-SKY conditions after preprocessing, ensuring balanced data length across comparisons. All preprocessing steps were applied uniformly to both intervention and control groups as part of the present secondary analysis.

### Feature Extraction

To comprehensively characterize SKY-related neural modulation, four complementary EEG feature representations were extracted independently from the preprocessed signals.

#### Raw EEG Statistical Features

Time-domain statistical descriptors were computed directly from the preprocessed EEG voltage signals. For each channel and epoch, measures including mean, variance, standard deviation, skewness, and kurtosis were extracted. These features capture baseline signal amplitude characteristics and overall neural variability associated with SKY-related neurophysiological state differences.

#### Short-Time Fourier Transform (STFT) Features

Spectral features were derived using the Short-Time Fourier Transform. For each EEG epoch, spectral power was computed and aggregated across time to obtain band-specific power estimates within canonical frequency bands (delta, theta, alpha, beta, and gamma). This approach captures frequency-domain characteristics informed by time-localized spectral analysis while producing fixed-length feature representations.

#### Discrete Wavelet Transform (DWT) Features

Multi-resolution analysis was performed using the Discrete Wavelet Transform to decompose EEG signals into approximation and detail sub-bands across multiple decomposition levels. For each EEG epoch, statistical summaries of the wavelet coefficients were computed for each sub-band, capturing frequency-specific neural activity at different temporal scales.

#### Coherence-Based Connectivity Features

Functional connectivity between EEG channels was quantified using magnitude-squared coherence computed for each electrode pair across canonical EEG frequency bands. Coherence values were averaged within each epoch to obtain static connectivity measures reflecting inter-regional synchronization and network-level neural coordination associated with SKY practice. These complementary feature representations capture distinct aspects of neural activity, including time-domain signal variability, spectral power dynamics, multi-resolution time–frequency structure, and functional connectivity. Each feature set was evaluated independently to assess its intrinsic discriminative capacity.

Feature importance for raw EEG voltage features was computed using impurity-based importance scores derived from the Random Forest classifier. Importance values were extracted within each fold of the leave-one-subject-out cross-validation (LOSO-CV) procedure and averaged across folds to obtain group-level scalp-level importance estimates.

### Machine Learning Models

To evaluate whether EEG-derived features could reliably distinguish pre- and post-intervention states, four machine learning models representing complementary learning paradigms were employed: Logistic Regression (LR), Support Vector Classifier (SVC), XGBoost, and a Multilayer Perceptron (MLP).

Logistic Regression was included as a linear baseline to assess the separability of the feature space under minimal model complexity. The Support Vector Classifier with a nonlinear kernel was used to capture margin-based nonlinear decision boundaries, which are commonly effective in high-dimensional EEG feature spaces.

XGBoost was selected as the representative tree-based ensemble model due to its ability to model complex nonlinear feature interactions through gradient-boosted decision trees while incorporating regularization mechanisms to mitigate overfitting. Given the conceptual overlap among ensemble tree-based methods, XGBoost was retained as a single representative, and Random Forest was not included in the final analysis.

Finally, a Multilayer Perceptron (MLP) was employed to assess the capacity of a feedforward neural network to learn nonlinear mappings from engineered EEG features without explicitly modeling temporal dependencies. Transformer-based architectures were excluded, as the feature extraction pipeline yielded fixed-length representations that no longer preserved the original temporal structure of the EEG signals, rendering such models theoretically mismatched to the data representation.

All models were trained and evaluated independently for each feature type (Raw EEG, STFT, DWT, and coherence) using a subject-independent Leave-One-Subject-Out Cross-Validation (LOSO-CV) framework.

### Model Training and Validation

Model performance was evaluated using a Leave-One-Subject-Out Cross-Validation (LOSO-CV) strategy. In each fold, all EEG epochs corresponding to a single subject were withheld as the test set, while epochs from the remaining subjects were used for model training. This procedure was repeated until each subject had served once as the held-out test participant.

The LOSO-CV framework ensured subject-independent evaluation, eliminated data leakage between training and testing phases, and provided a robust estimate of model generalization performance on unseen individuals.

### Performance Metrics

Model performance was evaluated using complementary classification metrics appropriate for a balanced binary classification setting. Accuracy was used to quantify the proportion of correctly classified EEG epochs, while the F1-score was reported to capture the balance between precision and recall. For each classifier and feature representation, mean performance values and variability across Leave-One-Subject-Out Cross-Validation folds were reported.

### Implementation Details

All analyses were implemented in Python using established scientific computing and machine learning libraries, including NumPy, SciPy, Pandas, Scikit-learn, and XGBoost. A consistent computational environment was maintained, and fixed random seeds were used to ensure reproducibility across all experiments.

## 3. Results

### Classification Performance Overview

Classification performance was evaluated using a Leave-One-Subject-Out Cross-Validation (LOSO-CV) framework to ensure subject-independent assessment. Results are reported separately for the intervention and control groups across four EEG feature representations: raw voltage features, Short-Time Fourier Transform (STFT) features, Discrete Wavelet Transform (DWT) features, and coherence-based connectivity features. Performance was quantified using accuracy and F1-score.

Across feature representations, the intervention group consistently demonstrated classification performance exceeding the 0.50 random-classification baseline, whereas the control group generally showed lower performance with greater variability. This indicates that the observed separability was specific to the SKY intervention and not attributable to general neural fluctuations or nonspecific effects.

### Raw EEG Voltage Features

Using raw EEG voltage features, moderate but consistent discrimination between pre- and post-intervention states was observed in the intervention group. Among the evaluated models, the Multilayer Perceptron (MLP) achieved the highest mean classification performance, reaching an accuracy of approximately 0.84 with a corresponding F1-score of 0.83 (Table 1). The Support Vector Classifier (SVC) and Logistic Regression demonstrated comparable but slightly lower performance, while XGBoost yielded marginally reduced accuracy and F1-score for this feature representation.

**Table 1.** LOSO-CV classification performance using raw EEG voltage features for the intervention group.

| Model | Accuracy | F1 Score | AUC-ROC |
| --- | --- | --- | --- |
| Multilayer Perceptron (MLP) | $0.840 \pm 0.137$ | $0.832 \pm 0.168$ | $0.938 \pm 0.121$ |
| Support Vector Classifier (SVC) | $0.814 \pm 0.156$ | $0.819 \pm 0.183$ | $0.938 \pm 0.160$ |
| Logistic Regression | $0.802 \pm 0.178$ | $0.802 \pm 0.217$ | $0.905 \pm 0.209$ |
| XGBoost | $0.781 \pm 0.177$ | $0.759 \pm 0.237$ | $0.905 \pm 0.183$ |

Overall, all classifiers achieved performance exceeding the 0.50 random-classification baseline in the intervention group, indicating that raw EEG voltage features contain detectable intervention-related information. However, the relatively higher variability across subjects suggests that these features provide moderate discriminative strength when used in isolation.

In contrast, control-group classification remained near the 0.50 chance level across all models (Table 2), with high inter-subject variability. No classifier showed consistent discriminative ability based on raw EEG voltage features alone.

**Table 2.** LOSO-CV Classification Performance Using Raw EEG Voltage Features (Control Group)

| Model | Accuracy | F1 Score | AUC-ROC |
| --- | --- | --- | --- |
| Multilayer Perceptron (MLP) | $0.463 \pm 0.282$ | $0.346 \pm 0.373$ | $0.472 \pm 0.365$ |
| Support Vector Classifier (SVC) | $0.572 \pm 0.186$ | $0.488 \pm 0.356$ | $0.453 \pm 0.425$ |
| Logistic Regression | $0.486 \pm 0.225$ | $0.389 \pm 0.351$ | $0.467 \pm 0.368$ |
| XGBoost | $0.420 \pm 0.254$ | $0.284 \pm 0.355$ | $0.298 \pm 0.366$ |

This lack of separability in the control group reduces the likelihood that

intervention-group performance reflects spurious structure or methodological bias, supporting its interpretation as intervention-related neural modulation.

The comparative performance of intervention and control groups across classifiers is illustrated in Figure 1.

**Figure 1.**
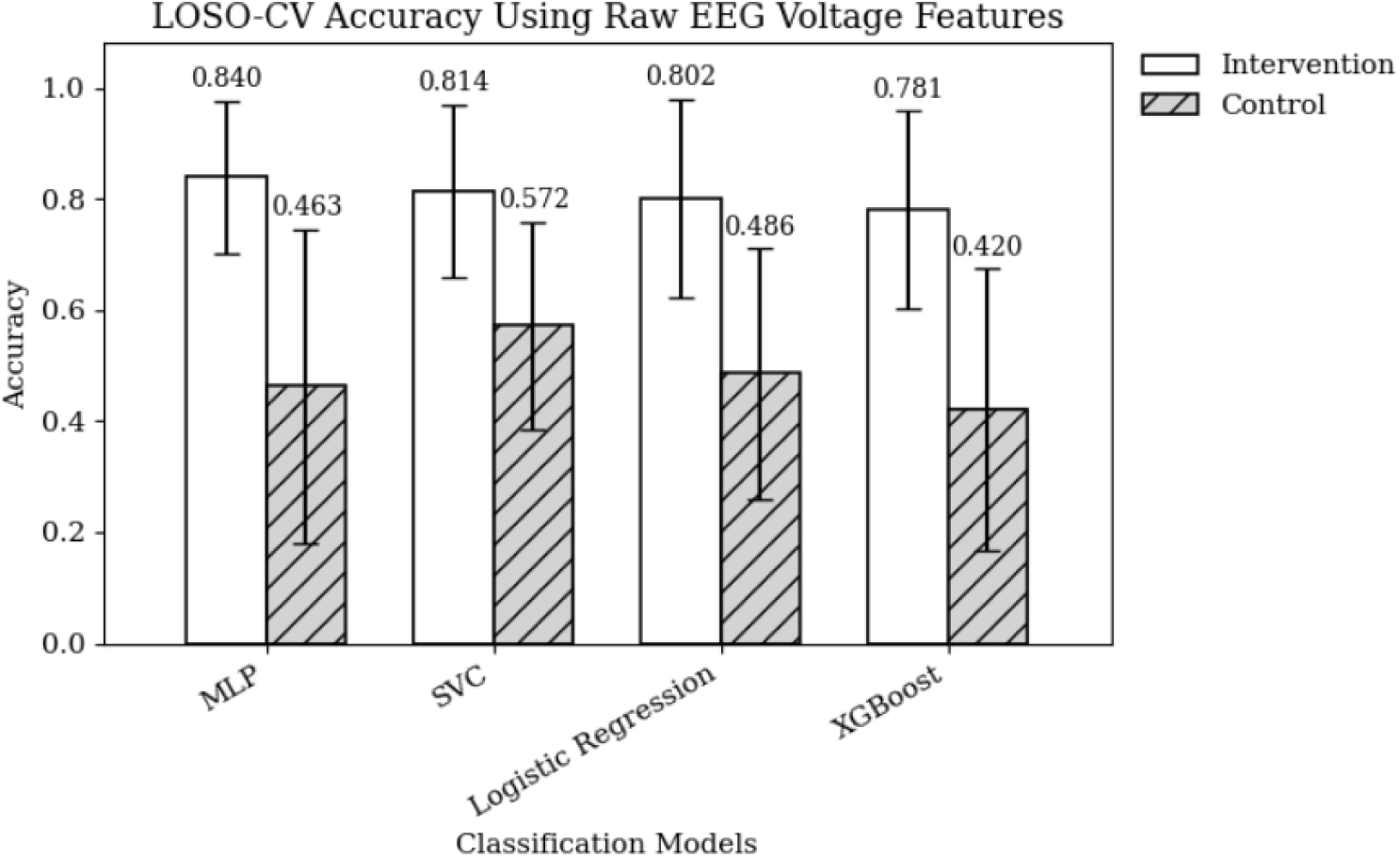
LOSO-CV classification accuracy using raw EEG voltage features for the intervention and control groups across four supervised machine learning models. The intervention group consistently achieved higher accuracy than the control group, a pattern consistent with intervention-related neural modulation.

#### Feature Importance and Scalp Distribution (Raw EEG Voltage)

Feature importance analysis of raw EEG voltage signals revealed distinct scalp-level patterns between the intervention and control groups (Supplementary Table S1).

In the intervention group, the most informative electrodes were predominantly located over frontal and frontal–midline regions (F3, Fz), with additional contributions from central (C4) and parietal–occipital (PO8) sites. This distribution indicates that intervention-related separability was driven primarily by anterior and central scalp activity, consistent with modulation of higher-order integrative and regulatory processes.

In contrast, the control group showed dominant contributions from posterior scalp regions, including parietal midline (Pz), bilateral parietal–occipital electrodes (PO3, PO8), and left temporal (T3) sites, with minimal frontal involvement. This pattern reflects greater inter-subject variability in posterior and temporal activity in the absence of a structured intervention, rather than systematic modulation.

These findings suggest that, even at the level of raw EEG voltage signals, the intervention condition exhibits a shift toward anterior–central scalp dominance, whereas control-group variability is primarily associated with posterior and temporal regions. Given the spatial limitations of scalp EEG and the influence of volume conduction, these observations are interpreted as scalp-level distributions rather than precise cortical sources.

These scalp-level patterns provided the rationale for further analysis using spectral, multi-resolution, and connectivity-based feature representations to better characterize the frequency-specific and network-level nature of the observed intervention effects.

### STFT-Based Spectral Features

STFT-derived spectral features substantially improved classification performance in the intervention group compared to raw EEG voltage features, indicating enhanced separability between pre- and post-intervention states in the spectral domain. As summarized in Table 3, all evaluated models achieved comparable mean accuracy, with the Multilayer Perceptron (MLP) yielding the highest mean accuracy (0.887). Notably, XGBoost exhibited markedly higher inter-subject variability (SD = 13.9%), whereas MLP, Logistic Regression, and Support Vector Classifier (SVC) demonstrated consistently low variability across LOSO folds, suggesting robust and stable discriminative patterns in the spectral domain.

**Table 3.** LOSO-CV classification performance using STFT features for the intervention group.

| Model | Accuracy | F1 Score | AUC–ROC |
| --- | --- | --- | --- |
| Multilayer Perceptron (MLP) | $0.887 \pm 0.035$ | $0.884 \pm 0.035$ | $0.981 \pm 0.012$ |
| XGBoost | $0.884 \pm 0.139$ | $0.882 \pm 0.159$ | $0.981 \pm 0.038$ |
| Logistic Regression (LR) | $0.875 \pm 0.038$ | $0.870 \pm 0.038$ | $0.970 \pm 0.017$ |
| Support Vector Classifier (SVC) | $0.846 \pm 0.051$ | $0.836 \pm 0.050$ | $0.946 \pm 0.020$ |

In contrast, classification performance in the control group was lower than in the intervention group but remained above the 0.50 chance baseline across models, with substantial variability across folds (Table 4). Although some models achieved moderate mean accuracy, the high inter-subject variability and less consistent F1 and AUC–ROC values indicate the absence of stable and robust discriminative patterns. This divergence between groups supports the interpretation that the stronger and more stable spectral separability observed in the intervention group is intervention-related rather than driven by spurious data structure.

**Table 4.** LOSO-CV classification performance using STFT features for the control group.

| Model | Accuracy | F1 Score | AUC-ROC |
| --- | --- | --- | --- |
| Support Vector Classifier (SVC) | $0.692 \pm 0.221$ | $0.670 \pm 0.317$ | $0.802 \pm 0.184$ |
| Logistic Regression (LR) | $0.615 \pm 0.143$ | $0.592 \pm 0.312$ | $0.712 \pm 0.222$ |
| XGBoost | $0.593 \pm 0.247$ | $0.484 \pm 0.363$ | $0.730 \pm 0.359$ |
| Multilayer Perceptron (MLP) | $0.598 \pm 0.164$ | $0.572 \pm 0.302$ | $0.695 \pm 0.256$ |

The comparative performance of intervention and control groups across classifiers is illustrated in Figure 2.

**Figure 2.**
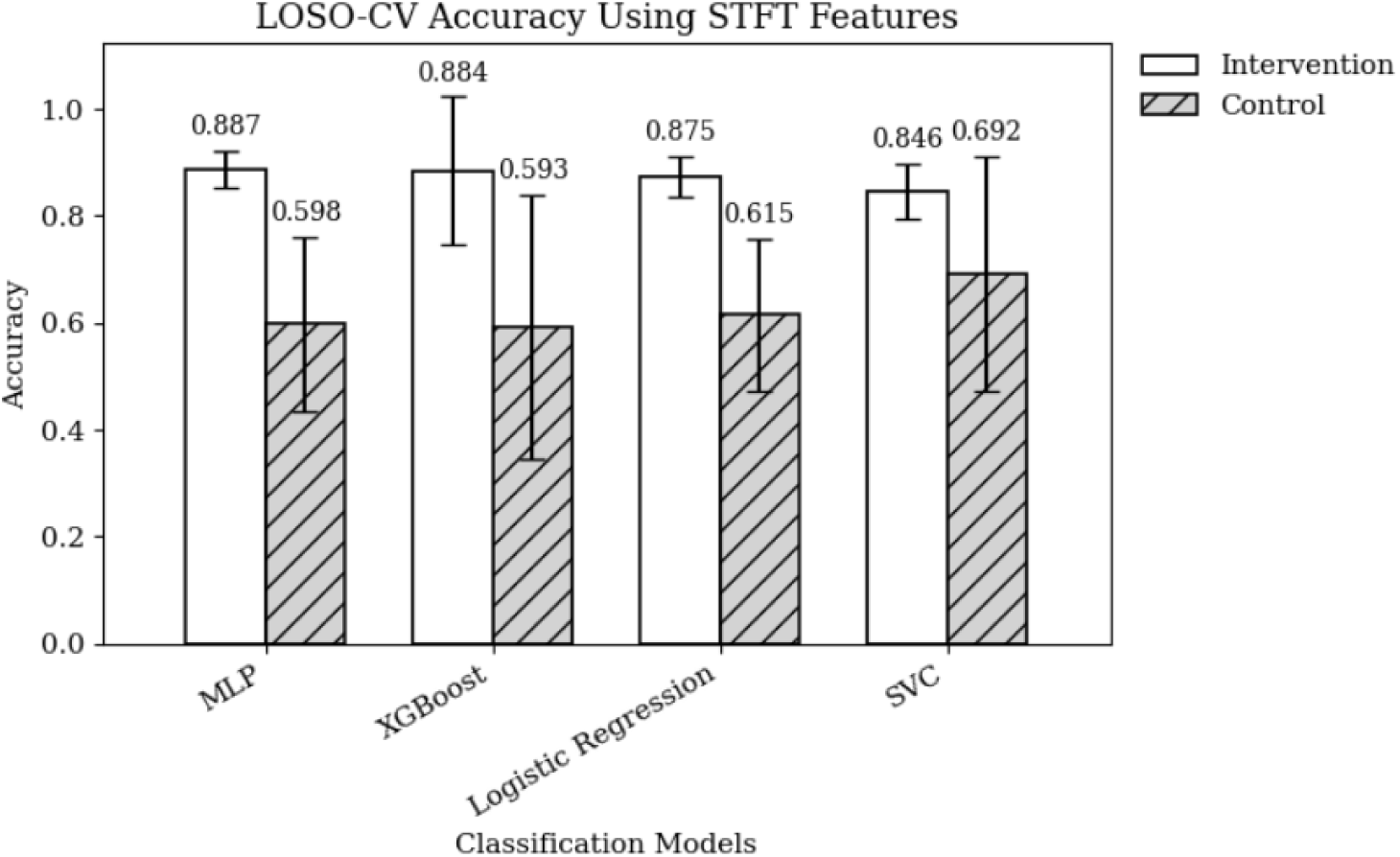
LOSO-CV classification accuracy using STFT features for the intervention and control groups across four supervised machine learning models. The intervention group consistently achieved higher accuracy than the control group, a pattern consistent with intervention-related neural modulation

#### Feature Importance and Scalp Distribution (STFT)

Intervention Group

**Table 5.** Top STFT features based on importance for the intervention group.

| Rank | Electrode (10–20) | Scalp Region | Mean Importance |
| --- | --- | --- | --- |
| 1 | Cz | Vertex | 0.0287 |
| 2 | C3 | Left motor | 0.0242 |
| 3 | Fz | Prefrontal | 0.0233 |
| 4 | Fz | Prefrontal | 0.0200 |
| 5 | Fz | Prefrontal | 0.0189 |

**Table 6.** Top STFT features based on importance for the control group.

| Rank | Electrode (10–20) | Scalp Region | Mean Importance |
| --- | --- | --- | --- |
| 1 | T7 | Left temporal | 0.0314 |
| 2 | O1 | Left occipital | 0.0302 |
| 3 | P7 | Left parietal | 0.0269 |
| 4 | O1 | Left occipital | 0.0223 |
| 5 | TP9 | Left temporo-parietal | 0.0176 |

**Table 7.** Band- and channel-level importance summary for the intervention group.

| Category | Top Contributor | Mean Importance |
| --- | --- | --- |
| Frequency Band | Gamma (relative) | 0.331 |
| Channel | Fz (Ch14) | 0.090 |

Feature importance analysis of STFT-derived features revealed that discriminative information was primarily localized to prefrontal and central scalp regions in the intervention group, with dominant contributions from gamma- and beta-band activity. As shown in Table 5, the most influential electrodes were located at the vertex and prefrontal midline (Cz, C3, and Fz), indicating strong contributions from central motor and prefrontal regions. Notably, Fz appears multiple times among the top-ranked features because it contributes across distinct frequency bands rather than representing duplicate entries.

At a summary level, Table 7 shows that relative gamma-band power emerged as the dominant frequency-band contributor, with Fz identified as the most informative channel. These findings indicate that intervention-related modulation was characterized by enhanced high-frequency oscillatory activity localized to anterior–central scalp regions. Importantly, multiple frequency bands (gamma, beta, and theta) contributed within the same prefrontal electrode (Fz), suggesting multi-scale spectral modulation rather than effects confined to a single oscillatory band.

In contrast, feature importance analysis for the control group revealed a predominantly posterior and temporal scalp distribution. As shown in Table 6, the most informative electrodes were located over left temporal and posterior regions (T7, O1, P7, and TP9), with no prominent involvement of prefrontal or central scalp areas.

Spectral contributions in the control group were distributed across posterior channels without a coherent spatial or frequency-band pattern, suggesting that these importance values reflect nonspecific inter-subject variability rather than systematic neural modulation, as summarized in Table 8.

**Table 8.**
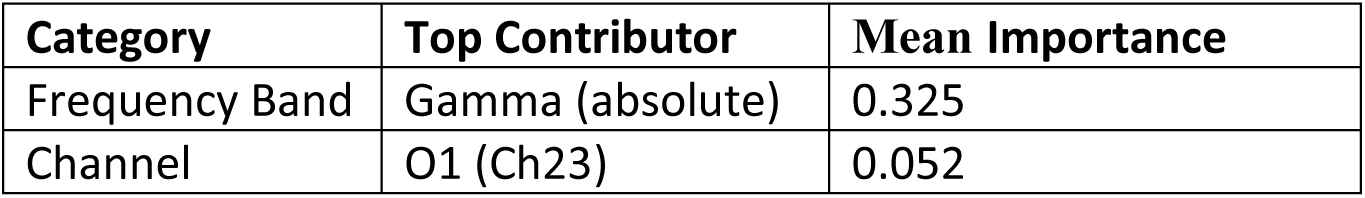
Band- and channel-level importance summary for the control group.

| Category | Top Contributor | Mean Importance |
| --- | --- | --- |
| Frequency Band | Gamma (absolute) | 0.325 |
| Channel | O1 (Ch23) | 0.052 |

Taken together, the contrast between anterior–central dominance in the intervention group and posterior–temporal dominance in the control group supports the specificity of STFT-based separability to the intervention condition.

As with all scalp-level analyses, these patterns reflect sensor-level distributions and should not be interpreted as direct cortical sources.

A complete list of STFT features and their importance values is provided in Supplementary Table S2.

### DWT-Based Multi-Resolution Features

DWT-based multi-resolution features demonstrated strong mean classification performance in the intervention group, with all models performing substantially above the 0.50 chance level under the LOSO-CV framework. The Multilayer Perceptron (MLP) achieved the highest mean accuracy (0.876) and F1-score, closely followed by Logistic Regression (0.859). Support Vector Classifier (SVC) and XGBoost also performed competitively, with AUC–ROC values exceeding 0.96 across models (Table 9).

**Table 9.** LOSO-CV classification performance using DWT features for the intervention group.

| **Rank** | **Model** | **Accuracy** | **F1 Score** | **AUC–ROC** |
| --- | --- | --- | --- | --- |
| 1 | Multilayer Perceptron (MLP) | $0.876 \pm 0.142$ | $0.867 \pm 0.185$ | $0.960 \pm 0.154$ |
| 2 | Logistic Regression (LR) | $0.859 \pm 0.150$ | $0.854 \pm 0.186$ | $0.932 \pm 0.181$ |
| 3 | Support Vector Classifier (SVC) | $0.837 \pm 0.158$ | $0.838 \pm 0.184$ | $0.977 \pm 0.050$ |
| 4 | XGBoost (XGB) | $0.831 \pm 0.170$ | $0.816 \pm 0.227$ | $0.976 \pm 0.050$ |

Despite high mean performance, DWT features exhibited more inter-subject variability than STFT features, with standard deviations ranging from approximately 14% to 17%, indicating that wavelet-based representations may be more sensitive to individual differences while retaining overall discriminative power (Table 9).

In the control group, performance remained near the 0.50 chance across all models, with variable F1 and AUC–ROC values, reflecting the absence of consistent discriminative patterns (Table 10).

**Table 10.**
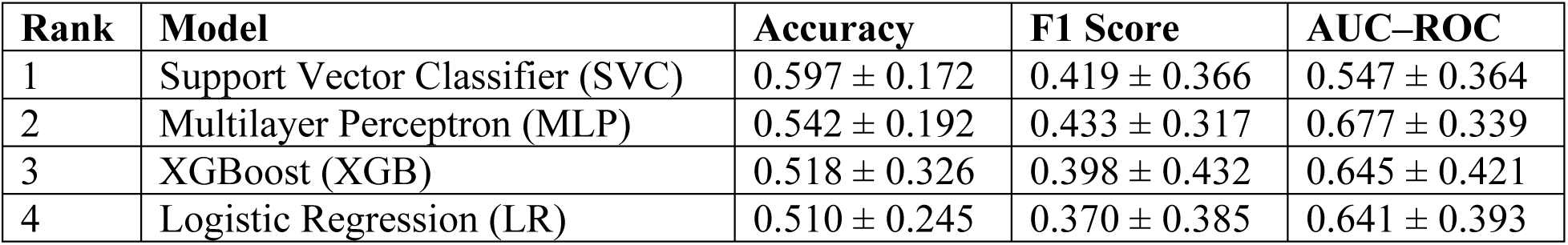
LOSO-CV classification performance using DWT features for the control group.

Overall, while DWT-based features offer high discriminative power on average, their increased inter-subject variability relative to STFT features suggests that multi-resolution wavelet representations may capture more subject-specific EEG signatures rather than uniformly consistent intervention-related patterns.

The comparative performance of intervention and control groups across classifiers is illustrated in Figure 3.

**Figure 3.**
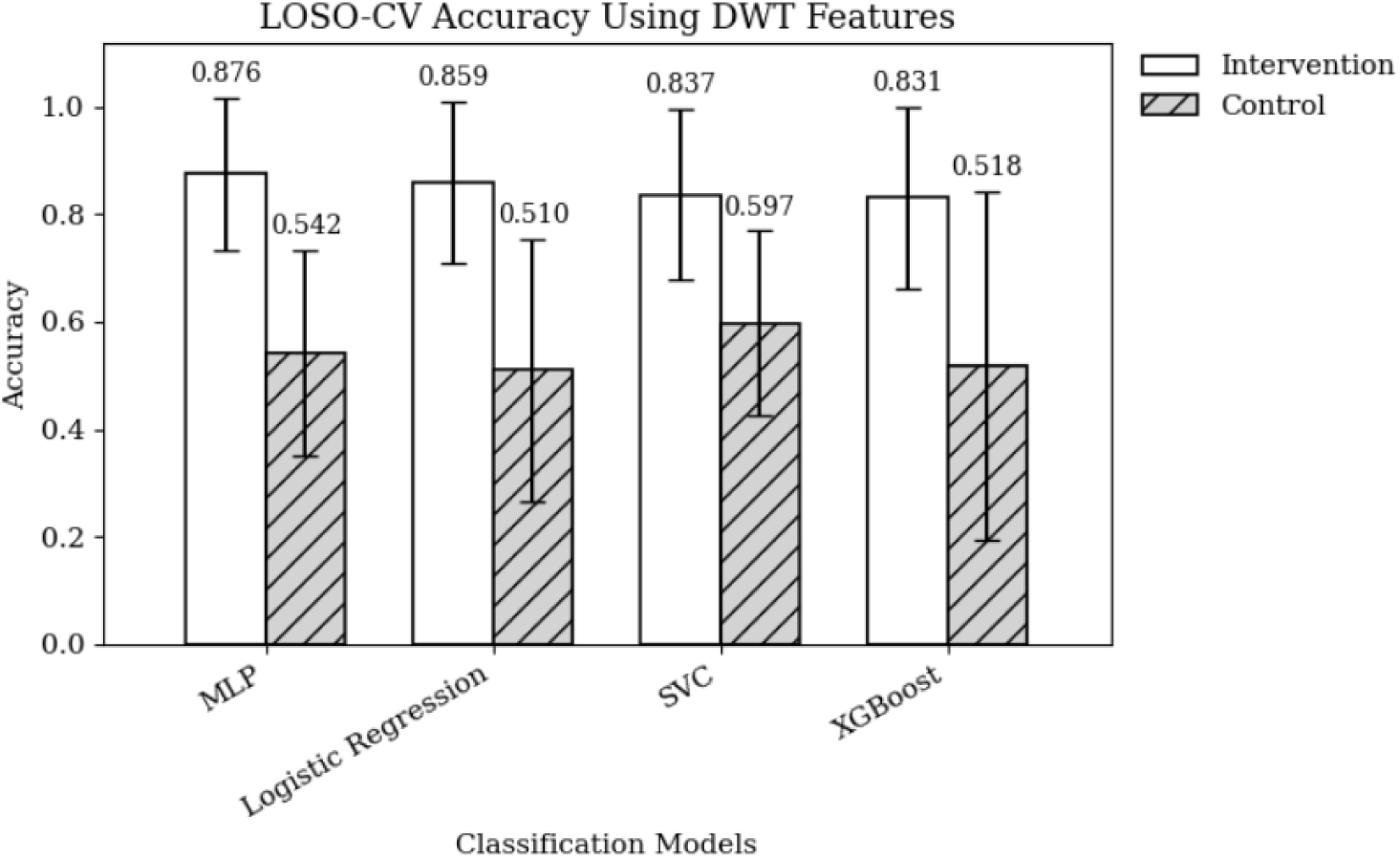
LOSO-CV classification accuracy using DWT features for the intervention and control groups across four supervised machine learning models. The intervention group consistently achieved higher accuracy than the control group, a pattern consistent with intervention-related neural modulation.

#### Feature Importance and Scalp Distribution (DWT)

Intervention Group

Control Group

Feature importance analysis of DWT-derived features revealed distinct scalp-level patterns between the intervention and control groups, consistent with the trends observed in raw voltage (Supplementary Table S1) and STFT-based analyses (Supplementary Table S2).

In the intervention group, the most informative features were predominantly associated with frontal–central scalp regions, with repeated contributions from FC5 (left frontal–central) across multiple frequency bands and temporal scales. Secondary involvement included temporal (T8) and parietal–occipital midline (POz) sites (Table 11). The repeated occurrence of FC5 reflects its relevance across multiple DWT components rather than a duplicate entry. This distribution suggests that intervention-related separability was driven primarily by anterior and fronto-central multi-resolution dynamics.

**Table 11.** Top DWT features based on importance for the intervention group.

| Rank | Electrode (10–20) | Brain Region | Mean Importance |
| --- | --- | --- | --- |
| 1 | FC5 | Frontal–Central (Left) | 0.0204 |
| 2 | POz | Parietal–Occipital (Midline) | 0.0186 |
| 3 | FC5 | Frontal–Central (Left) | 0.0179 |
| 4 | T8 | Temporal (Right) | 0.0171 |
| 5 | FC5 | Frontal–Central (Left) | 0.0155 |

In contrast, the control group exhibited strong posterior dominance, with the most influential features localized to parietal midline (Pz) and left parietal regions (P3, CP5) (Table 12). Similarly, the repeated presence of Pz reflects contributions across multiple scales rather than redundancy. Notably, frontal and frontal–central electrodes did not emerge among the top-ranked features in the control condition.

**Table 12.** Top DWT features based on importance for the control group.

| Rank | Electrode (10–20) | Brain Region | Mean Importance |
| --- | --- | --- | --- |
| 1 | Pz | Parietal (Midline) | 0.0263 |
| 2 | Pz | Parietal (Midline) | 0.0256 |
| 3 | P3 | Parietal (Left) | 0.0215 |
| 4 | CP5 | Parietal/Central (Left) | 0.0202 |
| 5 | Pz | Parietal (Midline) | 0.0192 |

These findings indicate that DWT-based features capture intervention-specific neural modulation characterized by fronto-central dominance, whereas control-group variability is primarily associated with posterior scalp regions. Given the multi-resolution nature of the DWT, this pattern suggests that the intervention induces changes not only in spectral content but also in the temporal structure of EEG activity across scales, particularly over anterior–central regions.

As with other feature representations, these results are interpreted at the level of scalp distribution rather than precise cortical localization, due to the spatial limitations of scalp EEG and the influence of volume conduction.

Across raw EEG voltage, STFT, and DWT feature representations, intervention-related separability consistently manifested as anterior–central (frontal and frontal–midline) dominance, with high-frequency activity (gamma and beta bands) contributing most prominently, whereas control-group discriminative patterns were primarily posterior–temporal, highlighting the specificity of SKY-induced neural modulation across multiple EEG feature domains.

### Coherence-Based Connectivity Features

Coherence-based connectivity features enabled robust discrimination between pre- and post-intervention states in the intervention group. Among the models, the Support Vector Classifier (SVC) achieved the highest mean accuracy and F1-score, indicating strong separability in the connectivity feature space. Multilayer Perceptron (MLP) and XGBoost showed comparable performance, suggesting that nonlinear decision boundaries effectively capture network-level EEG modulation. Logistic Regression performed above the 0.50 chance level, indicating some linear separability in coherence features. High AUC–ROC values across all classifiers further support the discriminative stability of coherence-based representations in the intervention group (Table 13).

**Table 13.** LOSO-CV classification performance using EEG coherence features for the intervention group.

| Model | Accuracy | F1 Score | AUC–ROC |
| --- | --- | --- | --- |
| SVC | $0.859 \pm 0.126$ | $0.857 \pm 0.149$ | $0.957 \pm 0.059$ |
| MLP | $0.849 \pm 0.126$ | $0.844 \pm 0.152$ | $0.939 \pm 0.081$ |
| XGBoost | $0.840 \pm 0.125$ | $0.826 \pm 0.169$ | $0.951 \pm 0.067$ |
| Logistic Regression | $0.805 \pm 0.127$ | $0.802 \pm 0.149$ | $0.889 \pm 0.110$ |

In the control group, classification performance remained near the 0.50 chance level across all models, with high inter-subject variability and no dominant classifier, consistent with the absence of systematic separable patterns (Table 14).

**Table 14.** LOSO-CV classification performance using EEG coherence features for the control group.

| Model | Accuracy | F1 Score | AUC-ROC |
| --- | --- | --- | --- |
| Logistic Regression | $0.661 \pm 0.196$ | $0.561 \pm 0.350$ | $0.795 \pm 0.244$ |
| XGBoost | $0.652 \pm 0.117$ | $0.546 \pm 0.281$ | $0.841 \pm 0.175$ |
| MLP | $0.642 \pm 0.170$ | $0.540 \pm 0.328$ | $0.779 \pm 0.236$ |
| SVC | $0.634 \pm 0.141$ | $0.548 \pm 0.328$ | $0.801 \pm 0.206$ |

The comparative performance of intervention and control groups across classifiers is illustrated in Figure 4.

**Figure 4.**
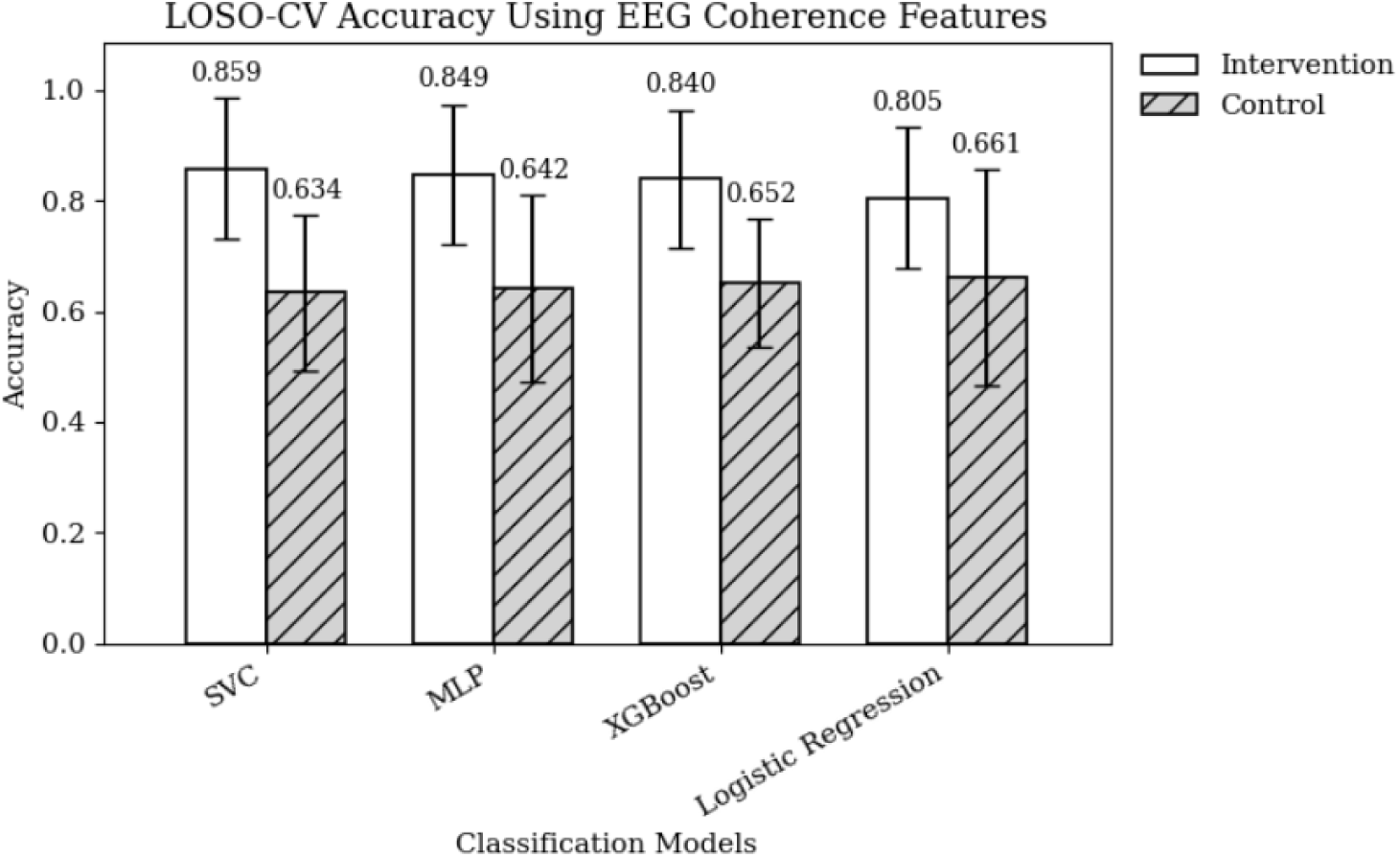
LOSO-CV classification accuracy using EEG coherence features for the intervention and control groups across four supervised machine learning models. The intervention group consistently achieved higher accuracy than the control group, a pattern consistent with intervention-related neural modulation.

#### Feature Importance and Network-Level Patterns (Coherence Features)

Intervention Group

The top 5 coherence features for the intervention group are shown in Table 15. Feature importance analysis of coherence-based connectivity features in the intervention group revealed that the most discriminative interactions were primarily concentrated between central and parietal regions (e.g., C4–Pz, CP6–POz). Individual feature importance values were relatively low, indicating that no single connection strongly dominated classification. Frequency contributions were mainly in the delta, theta, and beta bands, with minimal gamma involvement, suggesting that the intervention induced subtle, focused modulation of central–parietal networks.

**Table 15.** Top five coherence features based on importance for the intervention group.

| Rank | Coherence Feature | Electrode Pair & Region | Mean Importance |
| --- | --- | --- | --- |
| 1 | coh_delta_CP6_POz | CP6 (Parietal/Central, Right) – POz (Parietal/Occipital, Midline) | 0.00998 |
| 2 | coh_theta_C4_Pz | C4 (Central, Right) – Pz (Parietal, Midline) | 0.00965 |
| 3 | coh_delta_P8_FC2 | P8 (Parietal, Right) – FC2 (Frontal/Central, Right) | 0.00961 |
| 4 | coh_theta_C4_Cz | C4 (Central, Right) – Cz (Central, Midline) | 0.00925 |
| 5 | coh_delta_C4_Pz | C4 (Central, Right) – Pz (Parietal, Midline) | 0.00859 |

Control Group

The top 5 coherence features for the control group are shown in Table 16. In the control group, the top coherence features were more spatially dispersed, including central–occipital, frontal–temporal, and parietal–frontal interactions, with higher mean importance values than the intervention group. Contributions spanned delta to gamma bands, reflecting diffuse, nonspecific variability rather than structured intervention effects.

**Table 16.** Top five coherence features based on importance for the control group.

| Rank | Coherence Feature | Electrode Pair & Region | Mean Importance |
| --- | --- | --- | --- |
| 1 | coh_gamma_C4_Oz | Central (Right) – Occipital (Midline) | 0.01446 |
| 2 | coh_beta_F4_T7 | Frontal (Right) – Temporal (Left) | 0.01224 |
| 3 | coh_beta_C4_Oz | Central (Right) – Occipital (Midline) | 0.01065 |
| 4 | coh_beta_F4_F7 | Frontal (Right) – Frontal (Left) | 0.01058 |
| 5 | coh_beta_C4_FC5 | Central (Right) – Frontal–Central (Left) | 0.00975 |

In summary, coherence feature importance suggests that the intervention produced targeted, subtle network-level changes, while the control group exhibited broader, nonspecific patterns. The low absolute importance values highlight that, unlike raw, STFT, or DWT features, coherence changes were modest but regionally focused, supporting their role as complementary evidence of intervention-related neural modulation.

Across feature types, raw EEG, STFT, and DWT analyses revealed robust intervention-related separability with clear anterior–central dominance, while coherence features indicated subtler, spatially focused network interactions, collectively supporting the specificity of neural modulation induced by the intervention.

## 4. Discussion

This study examined the neurophysiological effects of SKY using multivariate EEG analysis and traditional machine learning across multiple feature representations, including raw EEG voltage, Short-Time Fourier Transform (STFT), Discrete Wavelet Transform (DWT), and coherence-based functional connectivity measures. Classification performance was evaluated using a leakage-free Leave-One-Subject-Out Cross-Validation (LOSO-CV) framework, enabling robust assessment of subject-level generalizability while minimizing contamination from non-neural sources. Across all feature types and classifiers, the intervention group consistently demonstrated higher classification accuracy and lower inter-subject variability than the control group, suggesting the presence of structured, intervention-related differences in EEG activity. These findings should be interpreted as evidence of discriminative patterns rather than direct causal neural changes.

A central finding was the consistent superiority of classification performance in the intervention group across all feature representations. This pattern suggests that SKY produces structured neural differences that enhance separability between pre- and post-intervention states, whereas the control group showed lower accuracy and greater heterogeneity. Notably, this divergence was evident not only in spectral features but also in connectivity-based representations, underscoring the robustness and specificity of the intervention effect.

The marked improvement in classification performance when moving from raw EEG features to spectral and connectivity-based representations underscores the greater sensitivity of features that capture oscillatory dynamics and inter-regional coordination. In particular, frequency-domain (STFT and DWT) and coherence-based features consistently outperformed raw voltage amplitudes, indicating that intervention-related neural modulation is more effectively expressed in spectral and network-level measures. This pattern is consistent with prior EEG studies of meditation and breathing-based practices, which report that intervention effects are more prominently reflected in frequency-specific activity and functional connectivity than in absolute signal amplitudes alone [9,10,13–15].

Raw EEG voltage features yielded only moderate classification performance, with variability-related statistics emerging as the most informative contributors. This pattern suggests that SKY influences the stability and dispersion of neural activity, potentially reflecting changes in global arousal or baseline neural regulation. However, raw voltage measures lack functional specificity and are more susceptible to inter-subject variability and residual noise, which likely contributes to their comparatively weaker discriminative power.

Spectral and time–frequency representations proved substantially more sensitive to intervention-related effects. STFT-based features achieved the highest peak performance in the intervention group, with the Multilayer Perceptron reaching approximately 89% accuracy under LOSO-CV. DWT-based features also demonstrated strong overall performance, indicating that SKY-related neural modulation spans multiple temporal scales. The convergence of findings across STFT and DWT analyses strengthens confidence that frequency-domain effects constitute a robust signature of the intervention rather than a representation-specific artifact. In contrast, both spectral approaches yielded performance near the 0.50 chance level and greater variability in the control group, reinforcing the specificity of the observed oscillatory changes to structured breathing and meditation.

Coherence-based connectivity features provided complementary evidence for network-level reorganization following SKY. In the intervention group, coherence-based models achieved high mean classification accuracy under the LOSO-CV framework, despite inter-subject variability that was comparable to that observed for raw EEG voltage features. This indicates that, although individual differences in connectivity patterns remain pronounced, coherence features nonetheless encode systematic intervention-related information at the group level.

Spectral and connectivity analyses revealed distinct but related patterns of neural modulation: while spectral features indicated increased beta-band power at specific scalp locations, coherence analysis showed reduced beta-band synchronization, consistent with a shift from globally coupled beta activity toward more localized processing. In contrast, theta-band activity exhibited consistent increases in both spectral power and coherence, accompanied by elevated delta-band coherence, suggesting enhanced low-frequency engagement and large-scale integrative dynamics. Similar frequency-specific modulation and synchronization effects have been reported in prior EEG studies of SKY and meditation practices [13–15].

Taken together, the results across raw EEG, spectral, and connectivity features provide a multi-level characterization of SKY-related neural modulation. All interpretations are made at the scalp level, and no claims are made regarding underlying cortical generators. Raw EEG features capture coarse shifts in signal variability, spectral representations reveal frequency-specific oscillatory patterns, and coherence features reflect large-scale coordination dynamics at the scalp level. The consistently higher classification performance observed in the intervention group across all feature representations indicates that SKY induces both localized and distributed neurophysiological effects. Importantly, the use of traditional machine learning models enabled objective, data-driven quantification of these effects without reliance on a priori assumptions regarding specific channels or frequency bands, extending prior EEG studies of SKY that have primarily relied on univariate statistical analyses [13,14].

## 5. Limitations and Future Directions

Several limitations warrant consideration. First, analyses were performed at the sensor level, which constrains anatomical interpretability and may be affected by volume conduction. Future studies incorporating source-space reconstruction could more precisely localize the cortical generators underlying the observed effects.

Second, the present work examined short-term pre–post effects following a single SKY session. Although prior studies have investigated longitudinal practice, most focus on durations up to three months [10–12]. Future research extending beyond this timeframe would be valuable for assessing the durability and stability of the observed neural changes.

Third, while multiple EEG feature representations were evaluated, cross-frequency interactions were not explicitly examined. Given the modulation of theta, beta, and delta bands observed here, future work could incorporate cross-frequency coupling measures to better characterize hierarchical neural dynamics.

Fourth, classical machine learning models were selected for their interpretability and robustness under limited sample sizes. More advanced temporal or representation-learning models may further enhance sensitivity, provided they are carefully validated to preserve generalizability.

Finally, the modest sample size—particularly in the control group—may limit statistical power and contribute to variability. Larger and more balanced cohorts would support more stable estimates and enable subgroup analyses.

Overall, addressing these limitations through source-resolved, longer-term, and methodologically enriched approaches will be important for advancing understanding of SKY-related neural mechanisms and supporting translation to stress and mental health applications.

---

**Supplementary Table S1.** Top Raw EEG Signal Feature Importance with Scalp Localization

Intervention Group
| Rank | Electrode (10–20) | Scalp Region | Mean Importance |
| --- | --- | --- | --- |
| 1 | F3 | Frontal (Left) | 0.0807 |
| 2 | P4 | Parietal (Right) | 0.0751 |
| 3 | FZ | Frontal (Midline) | 0.0685 |
| 4 | PO8 | Parietal–Occipital (Right) | 0.0539 |
| 5 | C4 | Central (Right) | 0.0451 |

Control Group
| Rank | Electrode (10–20) | Scalp Region | Mean Importance |
| --- | --- | --- | --- |
| 1 | PZ | Parietal (Midline) | 0.0737 |
| 2 | PO8 | Parietal–Occipital (Right) | 0.0605 |
| 3 | PO3 | Parietal–Occipital (Left) | 0.0515 |
| 4 | T3 | Temporal (Left) | 0.0466 |
| 5 | A1 | Reference / Auxiliary | 0.0449 |

**Supplementary Table S2.** Raw STFT Feature Importance with Scalp Localization

Intervention Group
| Rank | Raw Feature | Frequency Band | Channel | Electrode (10–20) | Scalp Region | Mean Importance |
| --- | --- | --- | --- | --- | --- | --- |
| 1 | gamma_rel_ch10 | Gamma | 10 | Cz | Central (vertex) | 0.0287 |
| 2 | beta_rel_ch11 | Beta | 11 | C3 | Left central (motor) | 0.0242 |
| 3 | beta_rel_ch14 | Beta | 14 | Fz | Prefrontal (midline) | 0.0233 |
| 4 | gamma_rel_ch14 | Gamma | 14 | Fz | Prefrontal (midline) | 0.0200 |
| 5 | theta_rel_ch14 | Theta | 14 | Fz | Prefrontal (midline) | 0.0189 |
| 6 | gamma_rel_ch18 | Gamma | 18 | C4 | Right central (motor) | 0.0186 |
| 7 | gamma_rel_ch13 | Gamma | 13 | F3 | Left frontal | 0.0186 |
| 8 | beta_rel_ch8 | Beta | 8 | FC1 | Left fronto-central | 0.0181 |
| 9 | gamma_rel_ch9 | Gamma | 9 | FC2 | Right fronto-central | 0.0172 |
| 10 | theta_rel_ch25 | Theta | 25 | Pz | Parietal (midline) | 0.0172 |
| 11 | gamma_rel_ch11 | Gamma | 11 | C3 | Left central (motor) | 0.0169 |
| 12 | gamma_rel_ch22 | Gamma | 22 | CP2 | Right centro-parietal | 0.0164 |
| 13 | beta_rel_ch13 | Beta | 13 | F3 | Left frontal | 0.0158 |
| 14 | beta_rel_ch15 | Beta | 15 | F4 | Right frontal | 0.0155 |
| 15 | gamma_rel_ch19 | Gamma | 19 | CP1 | Left centro-parietal | 0.0153 |

Control
| Rank | Feature | Frequency Band | Channel | Channel (10–20) | Scalp Region | Importance |
| --- | --- | --- | --- | --- | --- | --- |
| 1 | gamma_abs_ch5 | Gamma (absolute) | 5 | T7 | Left Temporal | 0.0314 |
| 2 | gamma_abs_ch23 | Gamma (absolute) | 23 | O1 | Left Occipital | 0.0302 |
| 3 | theta_rel_ch7 | Theta (relative) | 7 | P7 | Left Parietal | 0.0269 |
| 4 | theta_rel_ch23 | Theta (relative) | 23 | O1 | Left Occipital | 0.0223 |
| 5 | gamma_abs_ch4 | Gamma (absolute) | 4 | T8 | Right Temporal | 0.0176 |
| 6 | gamma_rel_ch29 | Gamma (relative) | 29 | TP9 | Left Temporo-Parietal | 0.0167 |
| 7 | theta_abs_ch7 | Theta (absolute) | 7 | P7 | Left Parietal | 0.0164 |
| 8 | gamma_rel_ch27 | Gamma (relative) | 27 | PO3 | Left Parietal–Occipital | 0.0157 |
| 9 | beta_rel_ch30 | Beta (relative) | 30 | PO4 | Right Parietal–Occipital | 0.0146 |
| 10 | gamma_rel_ch16 | Gamma (relative) | 16 | C4 | Right Central (Motor) | 0.0141 |
| 11 | gamma_abs_ch29 | Gamma (absolute) | 29 | TP9 | Left Temporo-Parietal | 0.0132 |
| 12 | theta_rel_ch17 | Theta (relative) | 17 | CP5 | Left Central–Parietal | 0.0113 |
| 13 | theta_rel_ch27 | Theta (relative) | 27 | PO3 | Left Parietal–Occipital | 0.0113 |
| 14 | gamma_rel_ch31 | Gamma (relative) | 31 | PO8 | Right Parietal–Occipital | 0.0110 |
| 15 | beta_rel_ch22 | Beta (relative) | 22 | Pz | Midline<br>Parietal | 0.0109 |

## Reference

[1] World Health Organization. Depression and Other Common Mental Disorders: Global Health Estimates. Geneva: WHO; 2017.

[2] Selye H. Stress and the general adaptation syndrome. Br Med J. 1950;1(4667):1383–1392.

[3] McEwen BS. Protective and damaging effects of stress mediators. N Engl J Med. 1998;338(3):171–179.

[4] Arnsten AFT. Stress signalling pathways that impair prefrontal cortex structure and function. Nat Rev Neurosci. 2009;10(6):410–422.

[5] Thayer JF, Lane RD. A model of neurovisceral integration in emotion regulation and dysregulation. J Affect Disord. 2000;61(3):201–216.

[6] Thayer JF, Åhs F, Fredrikson M, Sollers JJ, Wager TD. A meta-analysis of heart rate variability and neuroimaging studies. Biol Psychol. 2012;91(2):154–164.

[7] Sharma H, et al. Yoga and stress regulation mechanisms. Indian J Physiol Pharmacol. 2005;49(4):411–420.

[8] Porges SW. The polyvagal perspective. Biol Psychol. 2007;74(2):116–143.

[9] Streeter CC, et al. Effects of yoga on GABA levels. J Altern Complement Med. 2007;13(4):419–426.

[10] Brown RP, Gerbarg PL. Sudarshan Kriya yogic breathing in the treatment of stress, anxiety, and depression. J Altern Complement Med. 2005;11(4):711–717.

[11] Janakiramaiah N, Gangadhar BN, Murthy PJV, Harish MG, Subbakrishna DK, Vedamurthachar A. Therapeutic efficacy of Sudarshan Kriya Yoga in dysthymia: a randomized controlled trial. J Affect Disord. 2000;57(1–3):255–259.

[12] Zope SA, Zope RA. Sudarshan Kriya Yoga: breathing for health. Int J Yoga. 2013;6(1):4–10.

[13] Chandra S, et al. Effect of Sudarshan Kriya Yoga on EEG and ECG during multitasking. CognNeurodyn. 2017;11(4):371–385.

[14] Sharma R, Raj T, Juneja M. EEG-based classification before and after yoga and Sudarshan Kriya. Biomed Signal Process Control. 2019;52:121–129.

[15] Bhaskar L, Tripathi V, Kharya C, Kotabagi V, Bhatia M, Kochupillai V. High-frequency cerebral activation and interhemispheric synchronization following sudarshan kriya yoga as global brain rhythms: the state effects. Int J Yoga. 2020;13(2):130–136.

